# Unification of Countermanding and Perceptual Decision-making

**DOI:** 10.1101/158170

**Authors:** Paul G. Middlebrooks, Bram B. Zandbelt, Gordon D. Logan, Thomas J. Palmeri, Jeffrey D. Schall

## Abstract

Perceptual decision-making, studied using two-alternative forced-choice tasks, is explained by sequential sampling models of evidence accumulation, which correspond to the dynamics of neurons in sensorimotor structures of the brain^1 2^. Response inhibition, studied using stop-signal (countermanding) tasks, is explained by a race model of the initiation or canceling of a response, which correspond to the dynamics of neurons in sensorimotor structures^3 4^. Neither standard model accounts for performance of the other task. Sequential sampling models incorporate response initiation as an uninterrupted non-decision time parameter independent of task-related variables. The countermanding race model does not account for the choice process. Here we show with new behavioral, neural and computational results that perceptual decision making of varying difficulty can be countermanded with invariant efficiency, that single prefrontal neurons instantiate both evidence accumulation and response inhibition, and that an interactive race between two GO and one STOP stochastic accumulator fits countermanding choice behavior. Thus, perceptual decision-making and response control, previously regarded as distinct mechanisms, are actually aspects of more flexible behavior supported by a common neural and computational mechanism. The identification of this aspect of decision-making with response production clarifies the component processes of decision-making.

---

We investigated whether perceptual decisions and response inhibition, which have been studied with different tasks and explained by different models, can be performed concurrently, whether they are accomplished by different neurons in separate circuits or by a common pool of neurons in a single circuit and whether perceptual decision making and countermanding can be unified computationally.

Three macaque monkeys performed a visual saccade choice countermanding task. On each trial, a checkerboard was presented with varied coherence of cyan and magenta colors. The monkeys reported the majority color with a saccade to one of two peripheral visual targets (Fig. 1a). On a minority of trials a visual stop-signal was presented after a variable stop-signal delay. On no-stop trials reinforcement was earned for a correct choice. On stop trials reinforcement was earned for canceling the choice saccade. Performance data from each monkey replicated classic features of both tasks (Fig. 1b, Extended Data Fig. 1, Extended Data Table 1). Response times and error rate increased with difficulty across both task dimensions. The duration of response inhibition, known as stop-signal reaction time (SSRT), was invariant with coherence. These behavioral results, confirmed in humans^5^, show that perceptual decision making and response inhibition operate concurrently and efficiently instead of competing for common resources.

**Figure 1.**
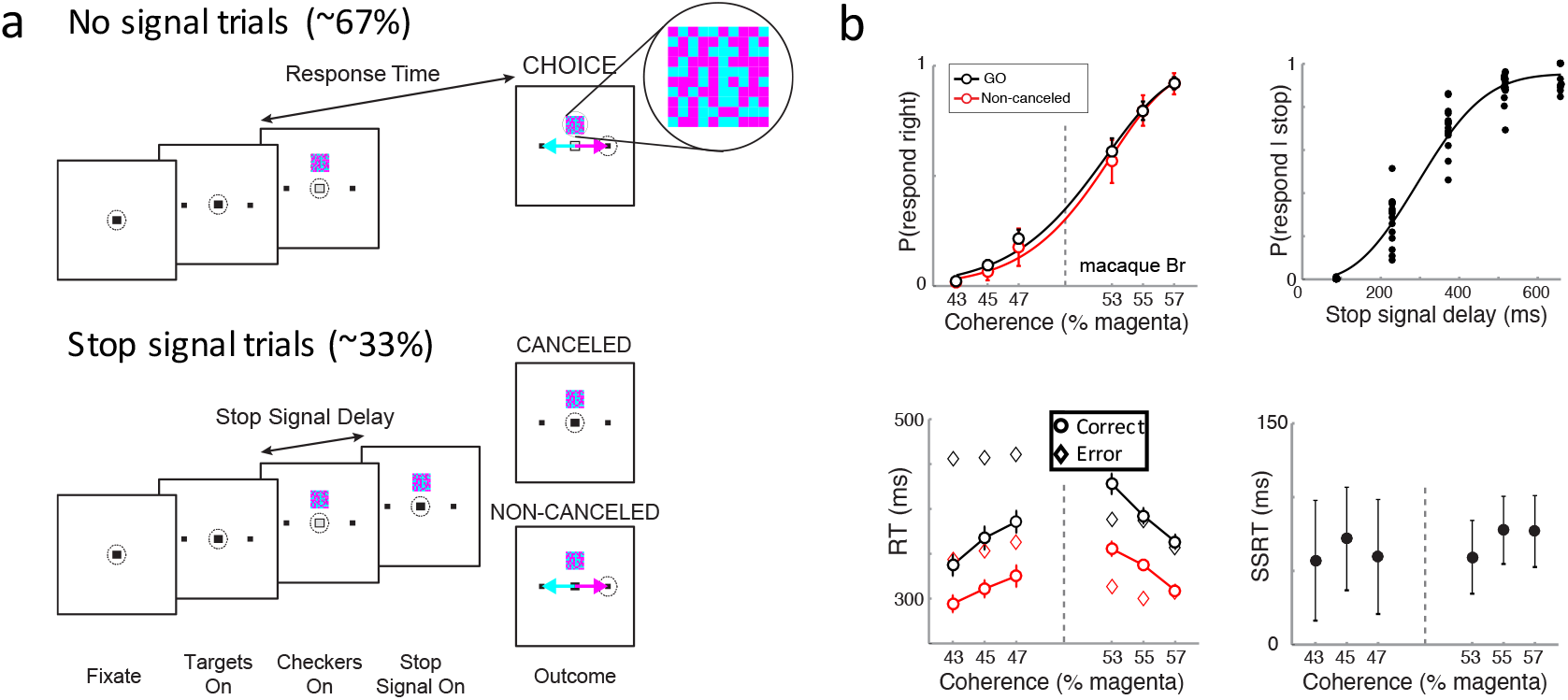
Perceptual decision countermanding task. **a**, No-stop trials (top panel) began by fixating a central spot. After a variable delay, two targets appeared in the periphery. After another variable delay, a 10 **x** 10 cyan-magenta checkerboard choice stimulus (magnified inset) appeared 3° directly above the fixation spot, and the fixation spot simultaneously disappeared. Fluid reward was delivered if monkeys shifted gaze to the target assigned to the respective colors. On a minority of trials (stop trials, bottom panel), the fixation spot reappeared after a variable stop-signal delay. Reward was delivered if monkeys canceled the planned saccade. b, Upper left, Psychometric functions for no stop (black) and non-canceled (red) trials. The variation of choice probability with color coherence is shown by the Weibull function fit to the mean (circles) and SD (bars). Upper right, Inhibition functions from all sessions. The variation of response inhibition stop-signal delay is shown by the Weibull function fit to the values. Lower left, Mean and SD of response time (RT) as a function of color coherence for correct (circle) and error (diamond) no-stop (black) and noncanceled (red) trials. Trends of correct RT highlighted by connecting lines. Noncanceled RT was systematically less than no-stop RT, justifying the application of the Logan race model. Lower right, Mean and SD stop-signal reaction time (SSRT) derived from race model as a function of color coherence. SSRT did not vary with decision-making difficulty.

In two monkeys, neural spiking was recorded in the frontal eye field (FEF), a prefrontal area at the interface of visual attention processing and saccade production^6^. This report is based on ~1400 neural spiking samples of which >1000 were modulated during the task with ~300 exhibiting presaccadic ramping activity. Of presaccadic units, ~60% were modulated with the difficulty of the perceptual choice in a manner that parallels the evidence accumulation process (Fig. 2, Extended Data Fig. 2), replicating previous observations^7^. Also, when choice saccades were canceled, ~40% of units modulated before SSRT in a manner sufficient to control saccade initiation, replicating previous observations^8 9 10^. We now report that ~25% of the presaccadic neurons exhibited both perceptual choice modulation and modulation before SSRT when saccades were canceled (Extended Data Table 2). Presaccadic ramping neurons in FEF have been identified computationally with the GO process of countermanding^11 12^ and with evidence accumulation for perceptual decision making^7 13 14^. These new results show that evidence accumulation and response inhibition are multiplexed in prefrontal cortex. If so, then in what sense can they be regarded as distinct processes?

**Figure 2.**
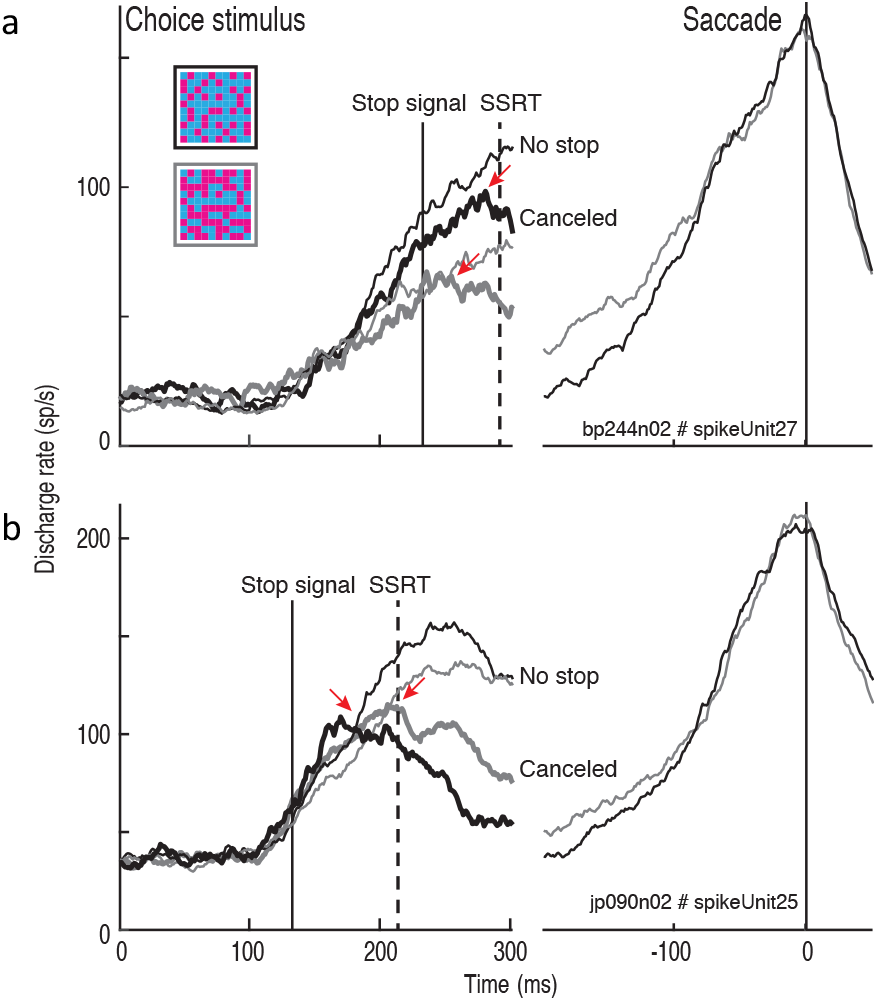
Neural mechanism of countermanding perceptual decision making. Representative neurons from monkey Br (**a**) and Jo (**b**) representing perceptual decision variable and instantiating GO process. Discharge rate is plotted as a function of time relative to choice stimulus presentation (left) and saccade (right). Activity accumulated faster with higher color coherence (thin solid lines: high color coherence is black; low color coherence is grey) but was inhibited before SSRT on canceled stop signal trials (thick lines).

Finding neural modulation for both perceptual decision difficulty and response inhibition in individual neurons entails a unity of choice and stopping models. To test that unity, we evaluated interactive race models consisting of 2 stochastic accumulators for each response alternative (GO_correct_, GO_error_) plus a STOP accumulator (Fig 3). We consider three mechanisms of choice (race, feed-forward inhibition, and lateral inhibition between GO units) and one mechanism of inhibition (inhibition from STOP to GO units). We tested five versions of parametric manipulation, allowing combinations of drift rate and/or starting point to vary across choice difficulty and response alternatives. We evaluated how well each version of the three model architectures fit the correct and error RT distributions, accuracy, and SSRT across levels of discrimination difficulty and stop-signal delays. Each architecture accounted for the combination of choosing and stopping performance (Fig 3, Table 3, Extended Data Table 3), with only modest differences in the goodness of fit of race, feedforward inhibition and lateral inhibition decision mechanisms. Decision difficulty was accounted for by variation in drift rates. Decision error rates were accounted for by combination of relative baseline levels and drift rates. Response inhibition was accounted for by late, potent inhibition of the GO units by the STOP unit. The trajectories of simulated GO and STOP units match the dynamics of observed neurons, producing a distribution of cancel times corresponding to that measured from the sampled neurons. These results demonstrate the graceful computational unification of the two primary models of decision-making and response control.

**Figure 3.**
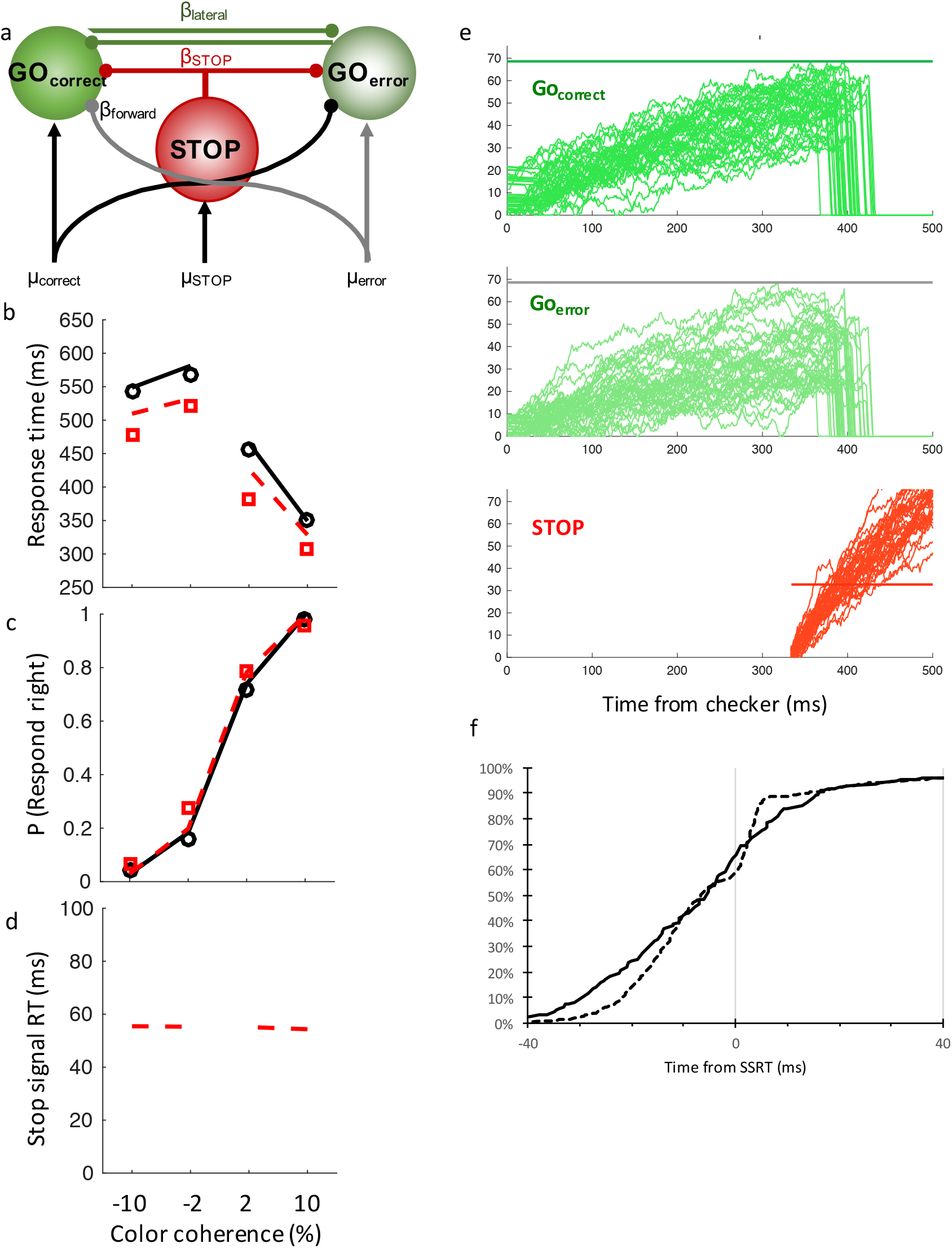
Simplest model of countermanding perceptual decision-making. **a**, Architecture of tested models. Alternative decisions are committed when the accumulated activation of one of the GO units reaches a threshold. The accumulation of each GO unit was driven either by the evidence supporting each alternative or by the difference in evidence through feedforward inhibition. The activation of each GO unit could simply race or be modulated through lateral inhibition. Decisions are canceled if the activation of a STOP process potently inhibits the GO units, **b,** Fit (lines) to correct (circle) and error (square) degenerate cumulative RT across color coherence. **c,** Fit to inhibition functions across color coherence. **d,** SSRT across color coherence. **e,** Representative trajectories of the GO_correct_, GO_error_, and STOP units. **f,** Observed (solid) and simulated (dashed) cancel times.

**Table 3.**
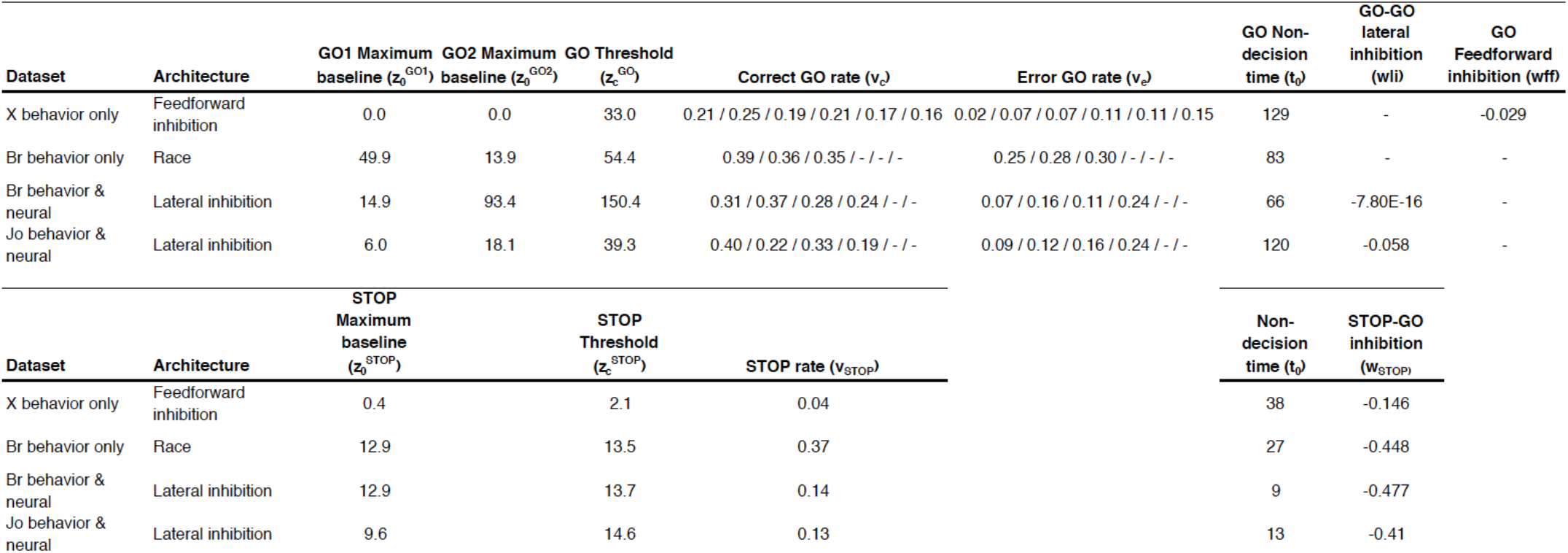
Parameters of best-fit models for each dataset. Each alternative choice architecture was endorsed by a dataset, but late-potent inhibition of the STOP unit was necessary to fit all datasets.

The performance measured during neural recordings corresponded to performance measured in other monkeys and four human participants^5^. Hence, while novel, the task demands were not unusually demanding. The patterns of neural modulation observed corresponded to patterns observed previously in monkeys performing tasks that require only saccade countermanding^8, 9^ or tasks that require only perceptual decision making^7 15 16^. Therefore, the novel observation of neurons that signal perceptual decisions being countermanded can be regarded as robust. Other neurons were found that only signaled perceptual decisions or that enacted countermanding. Further research is needed to elucidate the circuitry of these diverse neurons, but the incidence of neurons contributing to both functions confirms the unity of two previously distinguished cognitive operations.

The novel countermanding perceptual decision model fit performance of individual monkeys effectively as well as fits to perceptual decision or to countermanding performance alone^11 12 17 18^. Thus, the interactive race model unifies previously distinct model frameworks. The lateral inhibition and feedforward inhibition differences were less clear, but we regard this as evidence for the robustness of the unification and an indication that mathematically distinct but functionally similar neural networks can accomplish common functions^19^. Crucially, the necessity of late, potent inhibition of the STOP units on the GO units demonstrates that response inhibition does not operate as just another choice.

Our findings demonstrate a specific linkage between response preparation and perceptual decisions in prefrontal cortex that contradicts an earlier conclusion that frontal cortex does not contribute to evidence accumulation^20^ and can support performance without posterior cortical areas^21^. Under these testing conditions the decision process is identified with response choice, which corresponds grammatically and logical with *decide to*^22^. Thus, other neural processes must accomplish *decide that*, the categorization of observations that can be true/false, unlike choices. Evidence for neural processes of categorization distinct from response production has been reported in neurophysiological^23 24^, human EEG^25 26^ and functional imaging^27 28^ studies.

To conclude, new performance, neural and computational modeling results demonstrate that perceptual decisions and response inhibition are not separate processes but instead just different descriptions of a single process tested in different modes of operation.

## METHODS

### Monkey preparation

All experimental protocols were in accordance with National Institute of Health standards for care and use of laboratory animals and approved by the Vanderbilt University Institutional Animal Care and Use Committee. Performance data were collected from two macaques (1 female *M. mulatta* 5.4 kg identified as X, 15 sessions, and 1 male *M. radiata* 7.4 kg identified as *Br,* 15 sessions). Neural and performance data were collected from two macaques (Br, 82 sessions and another male *M. radiata* 11 kg identified as *Jo18* sessions). Cilux headposts and chambers (Crist Instruments) were implanted using standard surgical procedures. Monkeys sat comfortably with head restrained in a primate chair, facing a CRT (46° x 36° visual angle, 70 Hz refresh rate) in a dark room. Stimulus presentation, reward delivery, and task contingences were controlled by TEMPO/VIDEOSYNC software (Reflective Computing, Olympia, WA). Eye position was digitized at 1 kHz using an Eyelink 1000 eye tracker (SR Research) and streamed to a data acquisition system (Plexon). Fluid reinforcement was delivered with a solenoid-operated gravity-flow system.

### Perceptual decision countermanding task

The goal of the choice countermanding task was to choose whether a discriminatory stimulus contained more cyan or magenta and respond appropriately, though on some trials cancel the response when a stop-signal was presented. Each trial began when the subject fixated a spot in the center of the display (Fig. 1A: dashed circle is gaze position). After a variable duration (400-800 ms), two targets (1° square) appeared in the periphery, one in each hemifield 10° in amplitude from the central fixation spot and 180° from each other. Monkeys maintained fixation for another variable duration (400-800 ms), then the fixation spot was extinguished and simultaneously a choice stimulus appeared on the vertical meridian 3° above the central fixation spot. The choice stimulus was a 10x10 square checkerboard (magnified inset in Fig. 1) with a randomized pattern of isoluminant (30 cd/m^2^ on 13 cd/m^2^ gray background) cyan and magenta checker squares, and subtended 1°. The appearance of the choice stimulus and coincident disappearance of the fixation spot cued subjects to choose a saccade target by discriminating whether the checkerboard contained more cyan or magenta checkers. The assignment of cyan or magenta to leftward or rightward responses was counterbalanced across monkeys. Stimulus discriminability was manipulated trial to trial by randomly varying the color coherence of cyan and magenta checkers from among a set of 4 (in neural sessions) or 6 (in behavioral sessions) possible values. To obtain a broad range of choice accuracy, the coherence values were determined separately for each monkey (Br: [41%, 45%, 48%, 52%, 55%, 59%] or [42%, 47%, 53%, 58%]; Xe: [35%, 42%, 47%, 53%, 58%, 65%]; Jo: [40%, 44%, 56%, 60%]). After the choice stimulus appeared, on no-stop trials monkeys earned juice reward for a saccade to the correct target within 1000 ms. RTs were defined as the duration between the onset of the checkerboard stimulus and when eye movement velocity exceeded 30 degrees/sec away from fixation in the direction of one of the targets. A stop-signal was presented on ~33% of trials, but could vary between sessions (30-45% of trials) as we adjusted task parameters. During a stop trial the fixation spot reappeared after a variable stop-signal delay (SSD). Monkeys earned juice reward for canceling the saccade and maintaining gaze on the fixation spot. SSD was adjusted trial-to-trial using a staircase procedure within each color coherence level, to elicit equal proportions of canceled and noncanceled stop trials. SSD was increased randomly 1-3 steps (step size was constant within each session but could vary between sessions: [14, 29, 43, 57, 72, 86, or 100] ms/step = 3-7 70 Hz screen refreshes, using constant step increments within each session): after a canceled stop trial, and it was decreased randomly 1-3 steps after a noncanceled stop trial. The randomness of the stepping procedure was used to prevent SSD tracking by subjects, and SSD tracking was independent for each choice stimulus color coherence level. A saccade to the incorrect target on no-stop trials or a saccade to either target on stop trials resulted in a brief timeout. A saccade to the checkerboard stimulus at any time aborted the trial and resulted in a brief timeout. Aborts were rare (Br behavioral sessions: 1.6%; X behavioral sessions: 0.6%; Br neural recording sessions: 2.0%; Jo neural recording sessions: 1.6% of all trials). After receiving reward or timeouts, monkeys experienced a 1 second inter-trial interval. Macaque Br performed 15 behavioral sessions for modeling for a total of 33,258 trials, mean = 2,217 trials per session. Macaque Xe performed 15 behavioral sessions for modeling for a total of 25,927 trials, mean = 1,729 trials per session. Macaque Br performed 82 neural recording sessions for a total of 101,683 trials, mean = 1,240 trials per session. Macaque Jo performed 18 neural recording sessions for a total of 29,621 trials, mean = 1,646 trials per session. Of the neural recording sessions, 5 sessions each were chosen from Br and Jo for modeling. Modeled neural recording sessions were chosen based on behavioral performance and to ensure a nearly equal sample of trials from each monkey.

### Behavioral analyses

Each monkey provided multiple data collection sessions, so behavioral analyses throughout the study were performed within-subjects (across sessions). A few training sessions included unusually long noncanceled RTs during the earliest SSD trials. We consider these trials outliers, because they could be due to subjects not noticing the stop-signal (because it appeared so rapidly after fixation offset), or making an inadvertent saccade to a target without holding central fixation long enough to complete a stop trial. Therefore throughout our analyses we excluded trials that contained RTs greater or less than three standard deviations of the mean of the no-stop RT distribution in a given session. No results differed when the outlier trials were included. Choice performance was analyzed with respect to accuracy and RTs. Accuracy was quantified in psychometric functions, which plot the probability of making a saccade to the rightward target as a function of the percentage of rightward target color (*color coherence*) in the checkerboard. Psychometric functions were fit with a Weibull curve using maximum likelihood methods. The variation of RT was quantified as a function of color coherence in chronometric functions. To compare stopping performance across categorical choice difficulty, we estimated SSRT within each color coherence level. Given the large number of experimental conditions, we obtained at least ~50 stop-signal trials for each color coherence per session. Based on recent work, SSRT was calculated using the integration method (Verbruggen et al. 2013).

### Neural analyses

Neuronal spiking activity was recorded from two macaques (Br and Jo). Data were collected from Br over multiple sessions using various types of electrodes (47 sessions using single tungsten electrodes, 13 sessions using multi-channel (8 or 24) U-Probe vector arrays, and 19 sessions using 32-channel Neuronexus vector arrays. Neurons were considered task-modulated if their spiking activity varied as a function of visual stimuli and/or eye movements. 551 modulated units were recorded from Br. Data were collected from Jo over 18 sessions using 32-channel Neuronexus vector arrays. 376 modulated units were recorded from Jo. Each recording session, 800-1600 trials were collected to obtain enough data for statistical analyses. Electrode drift over time was unavoidable throughout each session. Thus single unit isolation often changed over the course of the session. Therefore we collapsed all units on each electrode channel into a single multi-unit for analyses, which provided stable spiking activity. Neurons were classified based on their response properties during task epochs.

We analyzed a population of neurons for this study consisting of two classes. Movement neurons modulated spiking activity leading to saccade initiation. Specifically, spike rates during the 50 ms leading to saccade exceeded those during the 300 ms prior to targets onset, assessed by paired t-test. Visuo-movement neurons modulated spiking activity the same as movement neurons, but in addition modulated 50-125ms after targets onset by paired t-test. We consider movement and visuomovement as one population in our analyses. Saccadic neuronal response fields were defined as the direction that elicited the highest spike rates during the 50ms prior to saccade initiation. Spiking activity was analyzed with respect to varying perceptual choice difficulty (color coherence) and response inhibition or initiation.

We tested whether unit spiking activity varied by color coherence in the interval following choice stimulus presentation leading to saccade initiation. For each neuron, the time when discharge rate began to represent the decision variable was determined based on a differential spike density function (SDF) test^8^. The mean no-stop trial SDF with the strongest color coherence signaling a response out of the movement field was subtracted from that of the strongest color coherence signaling a response into the movement field. A baseline difference discharge rate was calculated in the 500 ms prior to choice stimulus presentation. The beginning of decision variable representation was defined as the time at which the difference between mean SDFs reached 2 standard deviations of the baseline epoch differential SDF, and remained above 2 SD for at least 75 ms. A neuron contributed to the decision if it varied by both response direction and color coherence. To test whether a unit varied by response direction, we compared discharge rates between the choice stimulus and saccade initiation between nostop trials with responses into versus out of RF using a Wilcoxon rank sum test. To test whether a unit varied by color coherence, we compared mean SDFs during the decision epoch between easy and hard color coherence levels, within each response direction. Following previous work^7^, adapted for differences in behavioral tasks, a neuron contributed to the decision if the mean SDFs varied with color coherence for choices into the RF and either did not vary or varied in opposite sign for choices out of the RF.

To determine whether unit activity modulated with respect to response inhibition, we repeated the *cancel time* differential SDF test^8^. Within each condition (SSD **x** color coherence level), we subtracted mean SDFs of canceled stop trials SDFs from mean SDFs of latency matched no-stop trial, aligned on checkerboard onset. We defined a baseline epoch as 500 ms prior to checkerboard onset. Significant difference between mean SDFs was defined as the time at which the difference between mean canceled stop and latency-matched no-stop mean SDFs reached 2 standard deviations of the baseline epoch differential SDF, and remained 2 SD for at least 75ms. Cancel time was defined as the time of difference between the mean SDFs, minus the SSD within that condition, minus the SSRT within the session. We analyzed all stop trial conditions with at least 10 trials. A neuron contributed to response inhibition if it had a condition that canceled within 20 ms of SSRT for the session.

### Interactive race model

We extended the interactive race model (Boucher et al. 2007; Logan et al. 2015) to include perceptual two-alternative forced choice. The model included three stochastic accumulators. Two GO units accumulated evidence corresponding to the two decision alternatives. One STOP unit was activated upon stop-signal presentation. During no-stop-signal trials, a choice was made when one GO unit reached its threshold first. During stop-signal trials, if one of the GO units reached its respective threshold before the STOP unit, a noncanceled choice was made. If the Stop-signal reached its threshold, it inhibited the GO units. If the inhibition prevented both GO units from reaching threshold, the response was canceled.

We tested three alternative model architectures to determine which one provided the best fit to behavioral performance. The architectures differ with respect to how the GO units interacted with each other while accumulating toward threshold: 1) *Race:* units race to threshold independently, 2) *Lateral Inhibition:* units inhibited each other as a function of their respective level of activation 3) *Feedforward Inhibition:* the input to one unit inhibited the other. In each model the STOP unit inhibited both GO units uniformly via lateral inhibition.

Unit activation was governed by these equations:

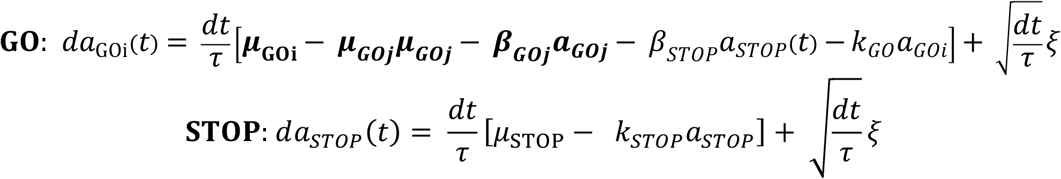

For each unit, the activation *a* increases as a function of multiple factors. *u* is the input to the respective unit (*drift rate*), *v* is the weight of feedforward inhibition (set to zero during Race and Lateral Inhibition model fits), *β* is the weight of lateral inhibition (set to zero for the GO unit during Race and Feedforward Inhibition model fits), *k* is a leakage parameter, and *ξ* is Gaussian noise. Other model parameters include unit threshold *z_c_* (discussed above for each unit); starting value *z_0_,* which is the initial level of unit activation, specified by a uniform distribution from trial to trial from a value of zero to z_0_; non-decision time (*t_nd_),* which is the time before a unit begins activation toward threshold, meant to account for duration of encoding and response execution by the brain.

For each architecture (Race, Lateral Inhibition, and Feedforward Inhibition), we fit 5 combinations of parameters free to vary across conditions: 1) Starting value varied between response directions (GO units); 2) Drift rate varied between choice difficulty (checker coherence), symmetric across response directions; 3) Starting value varied between response directions and drift rate varied between choice difficulty, symmetric across response directions; 4) Drift rate varied across choice difficulty, non-symmetric across response directions (so all 6 or 4 levels of choice difficulty, for the behavioral or neural datasets, respectively, had independent drift rates); 5) Starting value varied between response directions and drift rate varied between choice difficulty, non-symmetric across response directions. We chose to allow starting value to vary among response direction because each monkey displayed response time biases with respect to response direction. Allowing drift rate to vary among conditions emulates classic drift diffusion and stochastic accumulator models of choice. Models 1-2 were fit to test whether implausible models would fit the data well. Models 3-5 were fit to test whether plausible models would fit the data well, and better than models 1-2. Alternative parameter variation versions, allowing threshold and/or non-decision time to vary between response directions, did not improve fits to the behavioral performance. Models were fit by minimizing the chi-square statistic using the Nelder-Mead simplex algorithm in MATLAB. Model goodness of fit was assessed using the BIC statistic as follows:

Behavioral performance of monkeys Xe and Br were collected during sessions recording only behavior. Additional behavioral performance of Br and Jo was collected during neural recording sessions.

## Acknowledgements

This work was supported by F32EY023526, R01MH55806, R01-EY021833, P30EY008126, U54-HD083211, Vanderbilt Advanced Computing Center for Research and Education and Robin and Richard Patton through the E. Bronson Ingram Chair in Neuroscience. We thank J. Easley M. Feurtado, M. Maddox, S. Motorny, J. Parker, M. Schall, and L. Toy for animal care and other technical assistance. Correspondence concerning this article should be addressed to J. D. Schall, Vanderbilt Vision Research Center, Department of Psychology, Vanderbilt University, PMB 407817, 2301 Vanderbilt Place, Nashville, TN 37240-7817.

